# Alternative splicing regulation by GWAS risk loci for breast cancer

**DOI:** 10.1101/766394

**Authors:** Juliana Machado, Ramiro Magno, Joana M Xavier, Ana-Teresa Maia

## Abstract

Recent genome-wide association studies (GWAS) have revealed the association of hundreds of single nucleotide polymorphisms (SNPs) with breast cancer (BC) risk, which mostly locate in non-coding regions, suggesting regulatory roles to the causal variants. Functional characterisation of GWAS loci has been biased towards the effect of regulatory SNPs on transcription factor binding. Here we set out to determine the extent of the contribution of breast cancer risk-associated SNPs to alternative splicing (AS).

We screened genome-wide significant (*P* ≤ 5 × 10^*−*8^) BC risk SNPs for association with AS, using expression and genotype data from normal breast samples, from the GTEx project. We identified four splicing quantitative trait loci (sQTL). In locus 6p22.1, `rs6456883` is a significant cis-sQTL for the expression of `ZNF311` gene isoforms. Three SNPs in locus 8p23.3, `rs6682326`, `rs3008282` and `rs2906324`, were also identified as significant cis-sQTLs/svQTLs for the expression of *RPL23AP53* gene isoforms. In-silico functional analysis revealed that these variants can potentially alter enhancer splicing elements within the target genes.

Our work shows that BC risk-associated variants at two loci are associated with AS isoforms in normal breast tissue, thus demonstrating that AS plays an important role in breast cancer susceptibility. Furthermore, it supports that all cis-regulatory mechanisms should be considered in the functional characterisation of risk loci.

## Introduction

Recent genome-wide association studies (GWAS) have revealed the association of hundreds of single nucleotide polymorphisms (SNPs) with breast cancer (BC) risk, increasing the total knowledge of the risk’s genetic component to approximately 50%^1, 2^. However, GWASes do not identify directly the causative variants and are unable to pinpoint the underlying biological mechanisms. Interestingly, most risk-associated SNPs are located in non-coding regions, suggesting a regulatory role to these risk loci. Several studies have elucidated the biological mechanism of a small portion of risk loci (e.g.^3–5^), but have mainly focused on the role variants can have in regulating transcription via modulation of transcription factor binding to cis-regulatory elements (CREs). However, cis-regulation of gene expression can also be achieved by genetic variants altering binding of splicing factors^6–8^, including those associated with breast cancer risk^9, 10^, as well as miRNA binding^11, 12^. Our aim was to determine whether breast cancer risk-associated SNPs contribute to alternative splicing (AS) in normal breast tissue, the primary tissue of disease risk, and what the importance of this mechanism is to breast cancer risk.

## Results

### Splicing QTL analysis in breast cancer risk loci

Four hundred and thirty two BC GWAS SNPs with association p-value≤5 × 10^*−*8^, corresponding to 151 unique loci, were evaluated for their potential to modulate alternative splicing (Supplementary Table S1 indicates trait search criteria used). Upon identification of proxy SNPs, filtering of datasets and overlap of information, we obtained genotype data on 4,223 genetic variants and expression data on 120,812 transcripts, corresponding to 58,336 gene annotations. In total, sQTLseeker tested 50 gene-SNP pairs (Supplementary Table S2), and four significant sQTL associations were identified (FDR < 0.05) (Table 1 and Figure 1A and B).

**Table 1.**
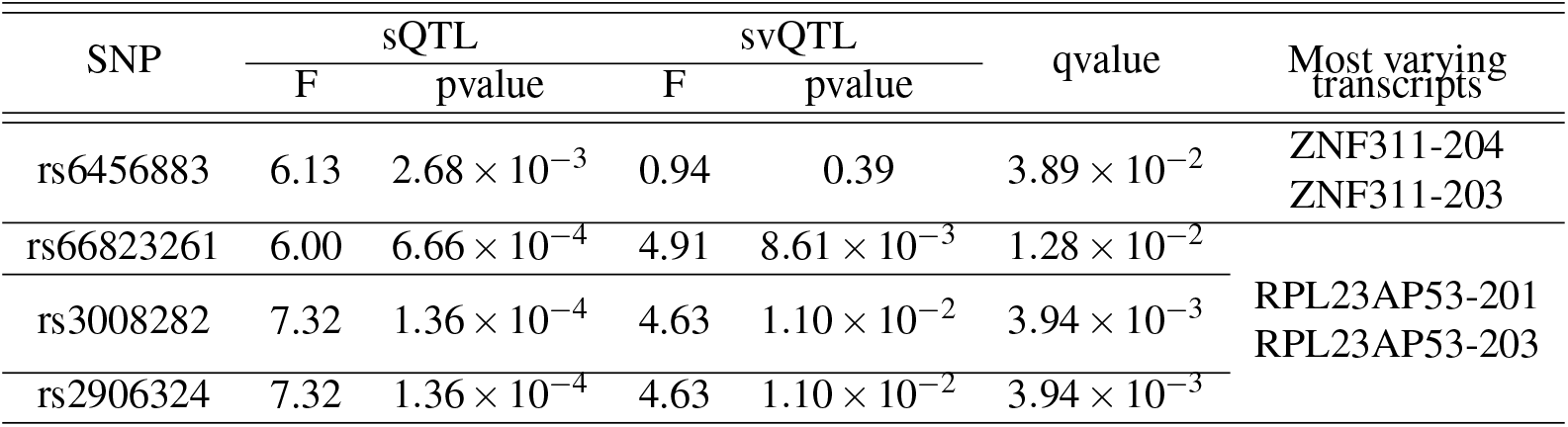
Significant sQTL associations for BC risk loci. Legend: F values and p-values, for both sQTL and svQTL, are indicated as obtained from sQTLseeker, and qvalues as obtained after multiple testing correction (5%FDR).

**Figure 1.**
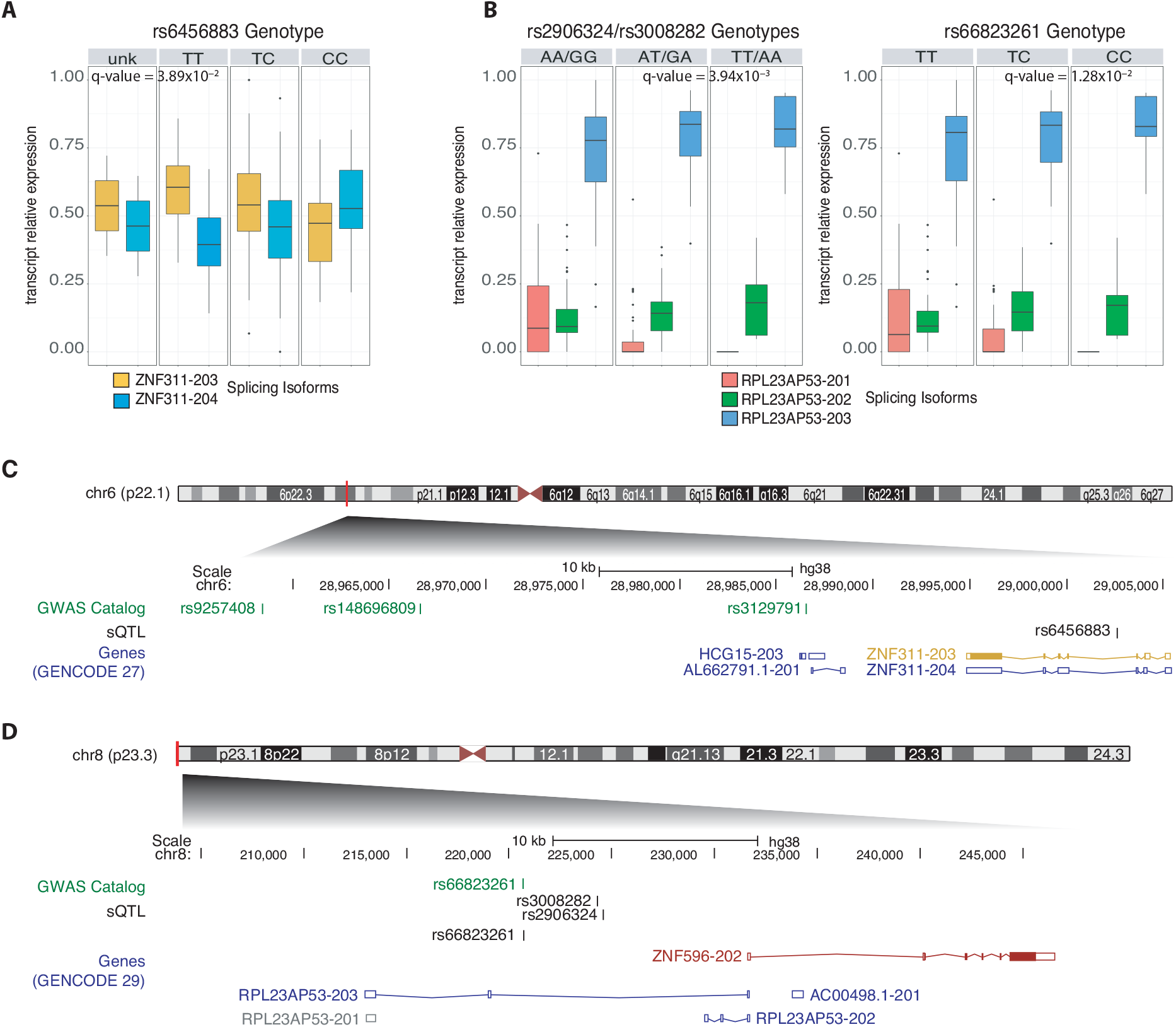
Breast cancer risk loci associated with alternative splicing. **A** and **B** - sQTLseekeR analysis for the risk variants in loci 6p22 (A) and 8p23 (B). Splicing isoforms for *ZNF311* and *RPL23AP53* are coloured differently as indicated below the graphs; y axis indicates the relative expression of the isoforms, i.e. the isoform specific RNA-seq read counts divided by the total read counts for the gene. Each panel represents a genotype group, as indicated in the grey bars above (unk - samples with unknown genotype). Each box-and-whiskers plot describes the distribution of expression values. The q-value of association is shown for each set. **C** and **D** - Schematic representation of breast cancer risk loci associated with alternative splicing (C - locus 6p22 and D - locus 8p23) with GWAS Catalog reported risk-associated variants displayed in green and identified sQTLs from current study displayed in black. Genes are displayed according to GENCODE 29 or 27, respectively, and colour-coded as follows: red for Ensemble protein coding genes; yellow for merged Ensembl/Havana protein coding genes; grey for non coding pseudogenes; and blue for non-coding processed transcripts.

rs6456883 was identified as a significant sQTL (q-value=3.89 × 10^*−*2^) for *ZNF311* (gene zinc finger protein 311 gene), localised in chromosome 6p22, and it localises to its third intron (Figure 1C). This variant is a proxy for the GWAS SNP rs9257408 (LD - *r*^2^ = 0.93) associated with breast cancer risk (C risk allele, p-val = 5 × 10^*−*8^, OR = 1.05 and 95%CI = [1.03-1.08])^1^ (Supplementary Table S3). In our study, the T allele of rs6456883, is associated with a predominance of the isoform *ZNF311-203*, the only protein producing transcript of *ZNF311* (Figure 1A). Conversely, the C allele is associated with predominance of the isoform *ZNF311-204*, an untranslated transcript characterised by the retention of an intron, resulting in a larger fourth exon. The same SNP is also an eQTL for the expression of *ZNF311* in testis, the tissue with higher expression of this gene, but not in breast tissue (data from^13^). We also found evidence of DAE for *ZNF311* in normal breast tissue (Supplementary Figure 1B), in two variants in LD with the sQTL variant (rs3129792 *r*^2^ = 0.74 and rs3129795 *r*^2^ = 0.28), further supporting the presence of cis-regulatory variants at this locus.

The other three significant sQTLs/svQTLs are rs66823261, rs3008282 and rs2906324 (q-values = 1.28 × 10^*−*2^, 3.94 × 10^*−*3^ and 3.94 × 10^*−*3^, respectively), associated with alternative splicing of the pseudogene *RPL23AP53* (ribosomal protein L23a pseudogene 53) (Figure 1B), located in chromosome 8p23, and widely expressed across tissues (data from^13^). These three SNPs are all located in the third intron of *RPL23AP53* (Figure 1D), with rs66823261 being the variant associated with risk to breast cancer in this locus (p-val = 3 × 10^*−*9^ for BC and 6 × 10^*−*9^ for ERnegBC, OR = 1.09 and 95%CI = [1.06-1.12])^14^ (Supplementary Table S3). The most predominant transcript *RPL23AP53-203*, and a shorter AS isoform containing only the first three exons (*RPL23AP53-202*), are both more expressed in the context of the minor allele of all three svQTLs. A minor AS isoform containing only a small portion of the 3’UTR of *RPL23AP53*, *RPL23AP53-201*, is the least expressed form, but more expressed in the context of the common allele. These three associations were significant for both sQTL and svQTL, meaning that there is heterogeneous variation between the levels of expression of all isoforms (heterogeneity of variation between the genotype groups). Due to this, we carried further analysis and found that rs3008282 and rs2906324 were significant eQTLs for each of the AS isoforms (Supplementary Figure 1A). We also found strong evidence of DAE for *RPL23AP53*, particularly at rs3008282 for which only one in 23 heterozygotes did not display a significant allelic expression difference (Supplementary Figure 1B). This result is a strong indicator that not only the expression of *RPL23AP53-203* is affected by cis-regulatory variants, but also that rs3008282 is a strong candidate causal variant of the DAE, as described previously^15^.

### In silico Predictive Functional Analysis

In order to unveil the possible functional impact of the significant sQTLs we investigated modification of donor/acceptor splice sites and ESE sites (exon splicing enhancer). We found a putative stronger intronic ESE site for *ZNF311* in the presence of the T allele of rs6456883, as predicted by two algorithms reported by the Human Splicing Finder tool (Supplementary Table S4). This is corroborated by the prediction using the NetGene2 website, which uses artificial neural networks applied to the prediction of splice site locations in human pre-mRNA (Supplementary Table S5). The allelic variation for the prediction is small (−0.86% for HSF matrix score and 33% for NetGene2 confidence), but it could explain the also small variation in the ratio of isoform expression. Nevertheless, we did not find evidence supporting that the rs6456883 alleles alter the binding of RNA-binding proteins.

For 8p23 locus we found predictions for all three sQTLs/svQTLs to affect both the binding of RBPs, as well as, potential ESE sites. Specifically, all three sQTL variants are predicted to potentially affect the binding of RBPs, with rs66823261 having the strongest supporting evidence (iCLIP and eQTL data)^16^ (Supplementary Table S6). The allelic differential predictions of RBP binding for these sQTLs, which could explain the association with different isoforms, are presented in Supplementary Tables S4, S7 and S8. rs66823261 and rs2906324 have the strongest predictions for the creation of intronic ESE sites, corroborated by the prediction of creation of new binding sites for several RBPs. Although rs3008282 is not predicted to modify AS by the Human Splicing Finder tool, RBPs are predicted to bind differentially to its alleles, with preferential binding to the A allele.

## Discussion

After more than a decade of GWAS efforts, the possible mechanisms by which cis-acting genetic variants regulate gene expression and contribute to disease susceptibility remain poorly understood. Most attempts at functionally characterising risk variants have focused on their potential to alter transcription factor binding. However, mRNA splicing is equally likely to be affected by genetic variants, whether by introducing changes in levels of expression and sequence of mRNA splicing-related genes or by modifying of RBP target sequences. The former has been assessed in liquid malignancies^17^, in lung cancer^18^, and even in breast cancer^10^. However, we believe that, in order to understand the biology of risk variants and how they increase risk during the lifetime of an individual, functional studies should be focused on the primary healthy tissue where disease develops. Thus, we carried an evaluation of the potential of GWAS risk variants to regulate AS in normal breast tissue.

Several computational tools are available that can identify cis-variants affecting alternative splicing^19^. In our study we used sQTLseekeR^20^, an R package which performs sQTL analysis by using the information of all alternatively spliced isoforms of a gene, and using their relative abundances as a vector, which is then used in a distance-based approach to test for association with genotype groups. Advantages of this package include the non-limitation of number of alternative splice isoforms that can be tested and the power to detect both cis- and trans-effects, as well as simple and complex alternative splicing events.

In our study, we identified four significant associations between BC risk-associated variants and differential transcript isoform expression levels and/or isoform relative amounts, from a total of 50 SNP-isoform pairs that were filtered for analysis by sQTLseekeR. Overall, this represents 8% of all 50 testable SNP-isoform pairs identified by sQTLseekeR, which suggest that AS is an important contributor to risk for BC. Additionally, none of these loci have previously been functionally characterised.

The significant association at 6p22 indicated that the intronic variant rs645883 is an sQTL for *ZNF311*, a gene encoding for a zinc-finger protein, predicted to be a transcription factor^21^, with broad expression across many tissues, including breast. This sQTL is in complete linkage disequilibrium with rs9257408, which is associated with risk to BC^1^, and also with lung cancer and tuberculosis^22, 23^. Also, in the GTEx^13^ database, rs6456883 is indicated as an eQTL for *ZNF311*, albeit only in testis, the tissue where it is most expressed.

In our study we found that the minor allele of rs6456883, is associated with decreasing relative expression of isoform *ZNF311-203*, a protein coding isoform, and increasing relative expression of isoform *ZNF311-204*, an unstranslated isoform with a retained intron. Intron retention is a known regulatory mechanism^24, 25^, which works in the way of decreasing the major isoform’s expression, by using the transcription machinery to produce a transcript which will not be translated. The in silico analysis suggests that rs6456883 contributes to a higher intron retention of *ZNF311* by weakening an intronic ESE. Altogether, our data suggests that lower expression of the main coding isoform might be the mechanism underlying risk to BC associated with this locus, but as the function of *ZNF311* remains unknown, we cannot predict more about its involvement in BC risk.

In locus 8p23, we detected significant svQTLs for three SNPs (rs66823261, rs3008282, rs2906324) associated with varying relative amounts of *RPL23AP53* AS isoforms. The *RPL23AP53* gene, is a processed pseudogene with unknown function, which is moderately expressed across all tissues (GTEx data^13^), and has three AS isoforms, with a predominant larger isoform (*RPL23AP53-203*). The locus has been associated with risk to breast cancer, in particular with risk to Estrogen Receptor Negative BC^14^. In fact, one of the identified svQTL signals corresponds to the GWAS SNP rs66823261. The other two SNPs, rs3008282 an rs2906324, are in almost complete LD with each other (*r*^2^ = 0.99), and in strong LD with the GWAS SNP (*r*^2^ = 0.82 and 0.83, respectively). All three svQTLs are located in the second intron of *RPL23AP53*, were characterised by increasing relative abundance of the main isoform associated with the minor alleles and have strong evidence of affecting the binding of RBPs. No sQTL is indicated in the GTEx database for *RPL23AP53*, but all three svQTLs we found have been identified as eQTLs for *RPL23AP53* in other tissues, but not in breast. Here, we found evidence for rs66823261 being an eQTL for *RPL23AP53* all three isoforms, and DAE data provides further supports the functional role of rs3008282 as a cis-regulatory SNP. Taken together, this data suggests that the risk for BC in this locus may be due to higher expression of the *RPL23AP53-203* isoform, whose expression is inversely correlated with that of *RPL23AP53-201*, the smallest isoform, consisting of a single exon. The higher expression of the pseudogene *RPL23AP53* could be working as a miRNA decoy protecting the parent gene Ribosomal Protein L23a from silencing, via competition for miRNA binding^26^, but the link to breast cancer increased risk needs further investigation. To our knowledge, this is the first study to systematically mine GWAS-significant SNPs for breast cancer to identify sQTL signals in normal breast tissue. We found that two risk loci, 6p22 and 8p23, may harbour causal variants that affect alternative splicing of two genes *ZNF311* and *RPL23AP53*, respectively, by altering the binding of RBPs and ESEs. This is the first functional analysis of these risk loci, with evidence supporting that the identified variants may be the causal of risk. Overall, our results demonstrate that alternative splicing is a mechanism by which risk-associated genetic variants, identified via GWAS, may be contributing to risk to breast cancer. Moreover, our work adds support to unbiased study approaches for complete unveiling of the biological mechanism underlying risk to breast cancer in particular, and complex diseases in general.

## Methods

### Data Sources

Breast cancer risk variants were retrieved, on 2018-06-12, from the GWAS Catalog website^27^, using a p-value cut-off significance of 5 × 10^*−*8^, and a list of related disease/trait with BC (Supplementary Table S1). Gene transcripts’ expression and genotyping data was obtained from GTEx project (dbGaP accession number: phs000424.v7.p2)^13^ on 2018-03-14. Data was retrieved for a total of 103 female breast samples. All data throughout the analysis was converted to the genome assembly GRCh38.

### Linkage disequilibrium (LD) analysis and Annotation

Variants in LD with the breast cancer GWAS SNPs were retrieved with the R package proxysnps (https://github.com/slowkow/proxysnps) using CEU population genotype data from the third phase of the 1000 Genomes Project^28^. Threshold parameters were set as *r*^2^ *≥* 0.8 and distance limit of *±*500 kb on either side of the queried SNP.

### Splicing QTL (sQTL) Analysis

Splicing QTL (sQTL) analysis was carried with the R package sQTLseekeR^20^, which filters data for a threshold of isoforms’ expression level of 0.01, threshold of gene expression of 0.01 and threshold of dispersion of 0.1. The authors differentiate between an sQTL, a genetic variant associated with changes in the splicing isoform ratios of a gene, and a splicing variation QTL (svQTL), a genetic variant associated with changes in the level of expression of a gene’s splicing isoforms, but not their relative amounts. Analysis was carried for both sQTLs and svQTLs, and p-values were corrected for multiple testing (FDR<5%).

### Expression QTL (eQTL) Analysis

The same data was used to perform eQTL analyses for the identified sQTL variants in the locus 8p23.3, as it was the only locus with evidence of eQTL in any tissue at the GTEx website. Analysis of variance (ANOVA) test was fitted, and p-values were corrected for multiple testing by using the Bonferroni method.

### Differential Allelic Expression (DAE) Analysis

Processed allelic expression and genotype data from normal breast tissue, described previously^29^, was used to determine differential allelic expression ratios (DAE) as log2 ((RNA allele A/RNA allele B)/ (DNA allele A / DNA allele B)). Analysis was carried in R using several Bioconductor packages^30^ and custom functions^31^. DAE was inferred when the absolute value of DAE was *≥* 0.58 (1.5 fold or greater difference between alleles).

### In silico Predictive Functional Analysis

In silico prediction of the effect of significant sQTL variants on the binding of RNA-binding proteins was carried using the RBP-Var database (RBP-VAR, http://www.rbp-var.biols.ac.cn/)^16^ and the RBPmap (http://rbpmap.technion.ac.il/)^32^ online tools. For splice variant and exonic splice enhancer analysis we used Human Splicing Finder (http://www.umd.be/HSF3/index.html)^33^ and NetGene2 Server (http://www.cbs.dtu.dk/services/NetGene2/)^34^ tools. In particular, we queried up to 200bp of sequence, centred on the sQTL variants, and searched for differential allelic predictions.

## Acknowledgements

Firstly, the authors would like to acknowledge the contributions made by all specimen donors who allow databases to be created and used to the good of medical care and science. This work was supported by national Portuguese funding through FCT-Fundação para a Ciência e a Tecnologia, UID/BIM/04773/2013-CBMR, SFRH/BPD/99502/2014 (JMX), and ALG-01-0145-FEDER-31477, which was co-funded by CRESC ALGARVE 2020. The authors would also like to thank the members of the Functional Genomics of Cancer group at CBMR for helpful discussions, Mr V. Morais at UAIC for administrative support, and Serv. de Informática, both at UAlg.

## Author contributions statement

J.M., R.M. and J.M.X. conducted bioinformatic analysis, A.T.M. conducted functional analysis and drafted the manuscript, R.M. and A.T.M. conceived and supervised the work. All authors reviewed the manuscript.

## Additional information

### Accession codes

Data used is available from GWAS Catalog (https://www.ebi.ac.uk/gwas/) and GTEx Project (https://gtexportal.org/home/) databases (dbGaP accession number: phs000424.v7.p2).

### Competing interests

The authors declare no competing interests.

