## Supplemental Material for "Alternative splicing regulation by GWAS risk loci for breast cancer"

### Supplementary Data

Juliana Machado, Ramiro Magno, Joana M Xavier and Ana-Teresa Maia *

April 2019

#### 1 Supplementary Tables

Table S1: GWAS Catalog Trait Filter Set

| **Disease/trait filter** |
|---|
| Breast Cancer |
| Breast cancer in BRCA2 mutation carriers |
| Breast cancer (estrogen-receptor negative, progesterone-receptor negative, and human epidermal growth factor-receptor negative) |
| Breast Cancer in BRCA1 mutation carriers |
| Breast cancer (estrogen-receptor negative) |
| Breast cancer (estrogen-receptor positive) |
| Cancer |
| Estrogen receptor status in HER2 negative breast cancer |
| Estrogen receptor status in HER2 positive breast cancer |
| Estrogen receptor status in breast cancer |
| HER2 status in breast cancer |
| Cancer (pleiotropy) |

Table S2: SNP/gene tested associations by sQTLseekeR before FDR correction.

| Gene | SNP | sQTL | | svQTL | | Most varying transcripts |
|---|---|---|---|---|---|---|
| | | F | pv | F | pv | |
| *ENSG00000119401.10* | 9_119457681_A_G_b37 | 0.4845450 | 0.6930564568 | 0.43591143 | 0.660295931 | *ENST00000373983.2*<br>*ENST00000450136.1* |
| *ENSG00000119401.10* | 9_119458931_G_A_b37 | 0.4544344 | 0.7157689812 | 0.44016405 | 0.656596794 | *ENST00000373983.2*<br>*ENST00000450136.1* |
| *ENSG00000184524.5* | 11_791462_G_A_b37 | 0.2711934 | 0.7740501213 | 0.75176011 | 0.487962963 | *ENST00000524587.1*<br>*ENST00000330106.4* |
| *ENSG00000197935.6* | 6_28958399_T_G_b37 | 0.2858527 | 0.6162318841 | 0.03574448 | 0.846392552 | *ENST00000377179.3*<br>*ENST00000483450.1* |
| *ENSG00000197935.6* | 6_28974265_G_T_b37 | 0.3175233 | 0.5942028986 | 0.03221894 | 0.856477890 | *ENST00000483450.1*<br>*ENST00000377179.3* |
| *ENSG00000197935.6* | 6_28977217_G_C_b37 | 0.2295420 | 0.6539130435 | 0.01094390 | 0.920868891 | *ENST00000377179.3*<br>*ENST00000483450.1* |
| *ENSG00000197935.6* | 6_28963248_T_G_b37 | 4.9275163 | 0.0081069865 | 0.77033516 | 0.460059910 | *ENST00000483450.1*<br>*ENST00000377179.3* |
| *ENSG00000197935.6* | 6_28964847_T_C_b37 | 4.9275163 | 0.0081069865 | 0.77033516 | 0.460059910 | *ENST00000483450.1*<br>*ENST00000377179.3* |
| *ENSG00000197935.6* | 6_28967038_A_G_b37 | 5.0101353 | 0.0075152943 | 0.72561745 | 0.483275087 | *ENST00000377179.3*<br>*ENST00000483450.1* |
| *ENSG00000197935.6* | 6_28968146_T_C_b37 | 4.9275163 | 0.0081069865 | 0.77033516 | 0.460059910 | *ENST00000483450.1*<br>*ENST00000377179.3* |
| *ENSG00000197935.6* | 6_28970369_T_C_b37 | 6.1323895 | 0.0026800177 | 0.94198925 | 0.392161757 | *ENST00000483450.1*<br>*ENST00000377179.3* |
| *ENSG00000197935.6* | 6_28970745_C_T_b37 | 3.7471055 | 0.0246716911 | 1.37217554 | 0.265601598 | *ENST00000377179.3*<br>*ENST00000483450.1* |
| *ENSG00000197935.6* | 6_28977925_C_T_b37 | 4.7842754 | 0.0092368241 | 0.61845456 | 0.546180729 | *ENST00000377179.3*<br>*ENST00000483450.1* |
| *ENSG00000223508.5* | 8_170692_T_C_b37 | 6.0009214 | 0.0006639329 | 4.90946297 | 0.008613009 | *ENST00000521854.1*<br>*ENST00000606975.1* |
| *ENSG00000223508.5* | 8_174284_A_T_b37 | 7.3183544 | 0.0001359863 | 4.62936133 | 0.010989011 | *ENST00000521854.1*<br>*ENST00000606975.1* |
| *ENSG00000223508.5* | 8_174546_G_A_b37 | 7.3183544 | 0.0001359863 | 4.62936133 | 0.010989011 | *ENST00000521854.1*<br>*ENST00000606975.1* |
| *ENSG00000258674.5* | 19_19625547_A_G_b37 | 2.3358608 | 0.0964754098 | 0.65829395 | 0.529180696 | *ENST00000555938.1*<br>*ENST00000586674.1* |
| *ENSG00000258674.5* | 19_19631444_A_T_b37 | 2.3358608 | 0.0964754098 | 0.65829395 | 0.529180696 | *ENST00000555938.1*<br>*ENST00000586674.1* |

Continued on next page

| Continuation of Table S3 | | | | | | |
|---|---|---|---|---|---|---|
| Gene | SNP | sQTL | | svQTL | | Most varying transcripts |
| | | F | pv | F | pv | |
| *ENSG00000258674.5* | 19_19631655_A_G_b37 | 2.3358608 | 0.0964754098 | 0.65829395 | 0.529180696 | *ENST00000555938.1*<br>*ENST00000586674.1* |
| *ENSG00000258674.5* | 19_19634863_C_CGCTGCGCCCAGGT_b37 | 2.3136892 | 0.0986065574 | 1.19645519 | 0.305836139 | *ENST00000555938.1*<br>*ENST00000586674.1* |
| *ENSG00000258674.5* | 19_19640524_T_C_b37 | 2.3358608 | 0.0964754098 | 0.65829395 | 0.529180696 | *ENST00000555938.1*<br>*ENST00000586674.1* |
| *ENSG00000258674.5* | 19_19643028_C_A_b37 | 2.3454187 | 0.0957377049 | 0.82898824 | 0.442760943 | *ENST00000555938.1*<br>*ENST00000586674.1* |
| *ENSG00000258674.5* | 19_19646272_T_C_b37 | 2.3454187 | 0.0957377049 | 0.82898824 | 0.442760943 | *ENST00000555938.1*<br>*ENST00000586674.1* |
| *ENSG00000258674.5* | 19_19648346_G_C_b37 | 2.3454187 | 0.0957377049 | 0.82898824 | 0.442760943 | *ENST00000555938.1*<br>*ENST00000586674.1* |
| *ENSG00000258674.5* | 19_19650096_A_G_b37 | 2.3454187 | 0.0957377049 | 0.82898824 | 0.442760943 | *ENST00000555938.1*<br>*ENST00000586674.1* |
| *ENSG00000258725.1* | 15_91512067_G_A_b37 | 1.3468704 | 0.2718736662 | 0.14387599 | 0.877777778 | *ENST00000554388.1*<br>*ENST00000556200.1* |
| *ENSG00000267058.1* | 19_44393257_A_G_b37 | 0.2972619 | 0.7329910141 | 2.02837234 | 0.139331970 | *ENST00000587128.1*<br>*ENST00000591815.1* |
| *ENSG00000267058.1* | 19_44393356_C_T_b37 | 0.3330024 | 0.7086007702 | 0.85376122 | 0.431629175 | *ENST00000591815.1*<br>*ENST00000587128.1* |
| *ENSG00000267058.1* | 19_44398536_CT_C_b37 | 0.2616146 | 0.7599486521 | 2.05163019 | 0.137286980 | *ENST00000587128.1*<br>*ENST00000591815.1* |
| *ENSG00000267058.1* | 19_44398940_A_G_b37 | 0.2685972 | 0.7567394095 | 0.86013593 | 0.427811861 | *ENST00000587128.1*<br>*ENST00000591815.1* |
| *ENSG00000267058.1* | 19_44399084_T_C_b37 | 0.2685972 | 0.7567394095 | 0.86013593 | 0.427811861 | *ENST00000587128.1*<br>*ENST00000591815.1* |
| *ENSG00000267058.1* | 19_44399565_C_T_b37 | 0.2685972 | 0.7567394095 | 0.86013593 | 0.427811861 | *ENST00000587128.1*<br>*ENST00000591815.1* |
| *ENSG00000267058.1* | 19_44400375_C_G_b37 | 0.2685972 | 0.7567394095 | 0.86013593 | 0.427811861 | *ENST00000587128.1*<br>*ENST00000591815.1* |
| *ENSG00000267058.1* | 19_44401793_G_A_b37 | 0.2685972 | 0.7567394095 | 0.86013593 | 0.427811861 | *ENST00000587128.1*<br>*ENST00000591815.1* |
| *ENSG00000267058.1* | 19_44402574_C_G_b37 | 0.2685972 | 0.7567394095 | 0.86013593 | 0.427811861 | *ENST00000587128.1*<br>*ENST00000591815.1* |
| *ENSG00000267058.1* | 19_44403055_G_A_b37 | 0.2685972 | 0.7567394095 | 0.86013593 | 0.427811861 | *ENST00000587128.1*<br>*ENST00000591815.1* |
| *ENSG00000267058.1* | 19_44404342_T_C_b37 | 0.2685972 | 0.7567394095 | 0.86013593 | 0.427811861 | *ENST00000587128.1*<br>*ENST00000591815.1* |
| *ENSG00000267058.1* | 19_44404398_G_T_b37 | 0.2384957 | 0.7785622593 | 0.88365680 | 0.417723245 | *ENST00000587128.1*<br>*ENST00000591815.1* |
| *ENSG00000267058.1* | 19_44404682_T_C_b37 | 0.2685972 | 0.7567394095 | 0.86013593 | 0.427811861 | *ENST00000587128.1*<br>*ENST00000591815.1* |
| *ENSG00000267058.1* | 19_44405044_A_T_b37 | 0.2685972 | 0.7567394095 | 0.86013593 | 0.427811861 | *ENST00000587128.1*<br>*ENST00000591815.1* |
| *ENSG00000267058.1* | 19_44405281_G_C_b37 | 0.2685972 | 0.7567394095 | 0.86013593 | 0.427811861 | *ENST00000587128.1*<br>*ENST00000591815.1* |
| *ENSG00000267058.1* | 19_44405287_C_T_b37 | 0.2685972 | 0.7567394095 | 0.86013593 | 0.427811861 | *ENST00000587128.1*<br>*ENST00000591815.1* |
| *ENSG00000267058.1* | 19_44406127_A_G_b37 | 0.3586215 | 0.6951219512 | 1.01926510 | 0.367007498 | *ENST00000587128.1*<br>*ENST00000591815.1* |
| *ENSG00000267058.1* | 19_44406291_T_C_b37 | 0.3586215 | 0.6951219512 | 1.01926510 | 0.367007498 | *ENST00000587128.1*<br>*ENST00000591815.1* |
| *ENSG00000267058.1* | 19_44406321_T_C_b37 | 0.3586215 | 0.6951219512 | 1.01926510 | 0.367007498 | *ENST00000587128.1*<br>*ENST00000591815.1* |
| *ENSG00000267058.1* | 19_44407487_A_G_b37 | 0.2908544 | 0.7374839538 | 0.95184944 | 0.391002045 | *ENST00000587128.1*<br>*ENST00000591815.1* |
| *ENSG00000267058.1* | 19_44408199_A_G_b37 | 0.2685972 | 0.7567394095 | 0.86013593 | 0.427811861 | *ENST00000587128.1*<br>*ENST00000591815.1* |
| *ENSG00000267058.1* | 19_44408753_T_C_b37 | 0.2685972 | 0.7567394095 | 0.86013593 | 0.427811861 | *ENST00000587128.1*<br>*ENST00000591815.1* |
| *ENSG00000267058.1* | 19_44408973_A_C_b37 | 0.2612866 | 0.7599486521 | 1.01333410 | 0.369188821 | *ENST00000587128.1*<br>*ENST00000591815.1* |
| *ENSG00000267058.1* | 19_44410846_A_G_b37 | 0.1436161 | 0.8620025674 | 0.90163080 | 0.410224949 | *ENST00000587128.1*<br>*ENST00000591815.1* |

Table S3: GWAS results from Milne *et al.* (2017) and Michailidou *et al.* (2017) for the significant sQTL at loci 6p22 and 8p23

| **SNP** | **pvalue** | **OR** | **95% CI** |
|---|---|---|---|
| rs9257408 | $5 \times 10^{-8}$ | 1.05 | [1.03-1.08] |
| rs66823261 | $3 \times 10^{-8}$<br>$6 \times 10^{-9}$ $(ER^{-})$ | 1.09 | [1.06-1.12] |

Legend: OR = per allele odds ratio, CI = Confidence Interval, ER = Estrogen Receptor.

Table S4: Human Splicing Finder Results

| **SNP** | **Splice Site Type** | **Motif** | **New Splice Site** | **Ref Allele** | **Alt Allele** | **Variation (%)** |
|---|---|---|---|---|---|---|
| rs64563883 | Acceptor | actatctcctaggt | accatctcctagGT | 81.74 | 81.04 | -0.86 |
| rs66823261 | Acceptor | aggatgacatagca | aggatgacacagCA | 65.98 | 73.67 | New Site 11.66 |
| rs2906324 | Acceptor | cagaatgtggaagc | cagaatgtggagGC | 41.32 | 70.26 | New Site 70.04 |

Table S5: NetGene2 Results

| Acceptor Splice Sites | Reference Allele | | | | Alternative Allele | | | |
|---|---|---|---|---|---|---|---|---|
| | **pos 5'-¿3'** | **phase strand** | **confidence** | **5' intron - exon 3'** | **pos 5'-¿3'** | **phase strand** | **confidence** | **5' intron - exon 3'** |
| | 510 | + | 0.33 | **T**ATCTCCTAG - GTTGCACCTC | 510 | + | 0.25 | **C**ATCTCCTAG - GTTGCACCTC |

Note: Search performed with 1001 bp sequence centred on rs6456883. Allele is shown in bold in intron-exon sequence.

Table S6: RBP-Var Results. Scores: 1e – Likely to affect RBP binding, RNA secondary structure and expression; 06 – Minimal possibility to affect RBP binding.

| SNP | CLIP binding | eQTL | Score |
|---|---|---|---|
| rs2906324 | Yes | No | 06 |
| rs3008282 | Yes | No | 06 |
| rs66823261 | Yes | Yes | 1e |

Table S7: Exonic splicing enhancers Results - ESEFinder

| SNP | Motif | Reference Allele | | Alternative Allele | |
|---|---|---|---|---|---|
| | | K-mer | Score | K-mer | Score |
| rs66823261 | SRSF1 (IgM-BRCA1) | | | CACAGCA | 3.55 |
| | SRSF5 | | | ACACAGC | 4.43 |
| rs3008282 | SRSF1 (IgM-BRCA1) | cagacta | 4.22 | | |
| | SRSF1 * | cagacta | 3.48 | | |
| | SRSF6 | tacaga | 3.74 | | |
| rs2906324 | SRSF2 | | | ggctgcag | 4.30 |

Table S8: RBP Map Results

| SNP | Motif | Reference Allele | | | Alternative Allele | | |
|---|---|---|---|---|---|---|---|
| | | K-mer | Z-Score | P-Value | K-mer | Z-Score | P-Value |
| rs66823261 | HNRNPL | | | | gacacag | 2.30 | $1.08 \times 10^{-2}$ |
| | | | | | ugacaca | 2.04 | $2.07 \times 10^{-2}$ |
| | | | | | acacagc | 3.18 | $7.29 \times 10^{-4}$ |
| | YBX1 | | | | cacagca | 3.27 | $5.32 \times 10^{-4}$ |
| | YBX2 | | | | cacagca | 3.03 | $1.23 \times 10^{-3}$ |
| rs3008282 | HNRNPL | acagacu | 1.92 | $2.74 \times 10^{-2}$ | | | |
| | RBM41 | | | | gucagu | 2.28 | $1.12 \times 10^{-2}$ |
| | RBMS1 | uacagac | 3.36 | $3.95 \times 10^{-4}$ | | | |
| | ZC3H14 | | | | ucuauuu | 2.13 | $1.64 \times 10^{-2}$ |
| rs2906324 | SRSF1 | uggaagc | 1.94 | $2.598 \times 10^{-2}$ | uggaggc | 2.13 | $1.65 \times 10^{-2}$ |
| | | | | | ggaggc | 1.71 | $4.36 \times 10^{-2}$ |
| | SRSF2 | | | | uggagg | 2.09 | $1.82 \times 10^{-2}$ |

### 2 Supplementary Figure

**Supplementary Figure 1**

A - eQTL analysis from GTex project data for sQTL variants identified in the 6p23.3 locus. p-values correspond to analysis of variance (ANOVA) tests, implemented in R. Adjusted p-values were corrected for multiple testing using the Bonferroni method.

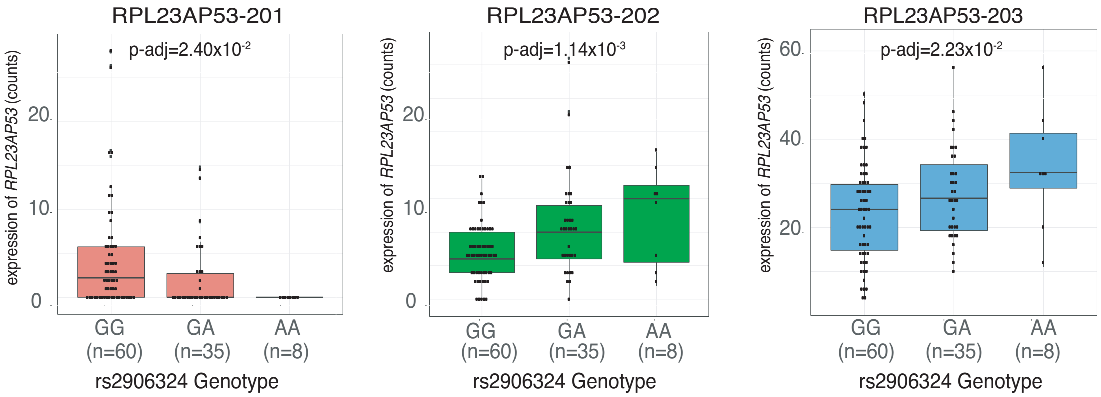


B - Differential alllelic expression analysis for *ZNF311* and *RPL23AP53* genes. Each dot is an heterozygous individual for the SNPs indicated in the x-axis. DAE ratios are calculated as $\log_2$(expression of allele A/ expression of allel B). Dashed lines correspond to DAE = 0.58 and -0.58, with samples above and below, respectively, considered to display significant allelic differences in expression (greater than 1.5 fold of either allele).

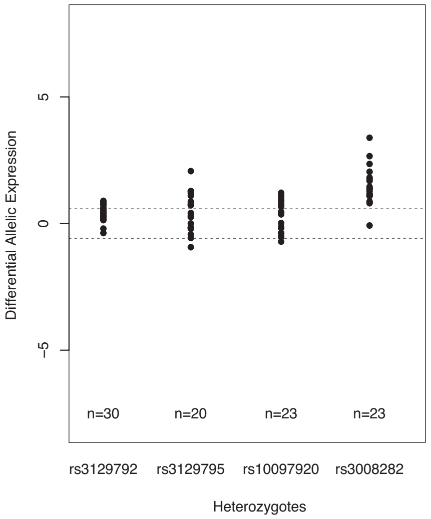
